# Layering genetic circuits to build a single cell, bacterial half adder

**DOI:** 10.1101/019257

**Authors:** Adison Wong, Huijuan Wang, Chueh Loo Poh, Richard I Kitney

**Author notes:** To whom correspondence should be addressed: Division of Bioengineering, School of Chemical and Biomedical Engineering, Nanyang Technological University, Singapore 637459, SG. Correspondence may also be addressed to: Centre for Synthetic Biology and Innovation, and Department of Bioengineering, Imperial College London, London SW7 2AZ, UK. Present Address: NUS Synthetic Biology for Clinical and Technological Innovation, National University of Singapore, Singapore 117456, SG.

## Abstract

Gene regulation in biological systems is impacted by the cellular and genetic context-dependent effects of the biological parts which comprise the circuit. Here, we have sought to elucidate the limitations of engineering biology from an architectural point of view, with the aim of compiling a set of engineering solutions for overcoming failure modes during the development of complex, synthetic genetic circuits. Using a synthetic biology approach that is supported by computational modelling and rigorous characterisation, AND, OR and NOT biological logic gates were layered in both parallel and serial arrangements to generate a repertoire of Boolean operations that include NIMPLY, XOR, half adder and half subtractor logics in single cell. Subsequent evaluation of these near-digital biological systems revealed critical design pitfalls that triggered genetic context dependent effects, including 5’ UTR interference and uncontrolled switch-on behaviour of σ54 promoter. Importantly, this work provides a representative case study to the debugging of genetic context dependent effects through principles elucidated herein, thereby providing a rational design framework to program single prokaryotic cell with diversified digital operations.

## INTRODUCTION

Gene regulation in biological systems behaves like molecular computers whereby the gene’s output can be modelled as on-off states of Boolean (digital) logic [1-3]. However, programming gene regulation is far from trivial and requires considerable time and effort during functional testing and tuning of the synthetic genetic circuits under development. Apart from the scarcity of reliable and well characterised biological parts, digital performance in biological systems is further impacted by the cellular and genetic context dependent effects of the biological parts which comprise the circuit [4-6]. Recent studies have shown that genetic cross-talks between the engineered circuits and endogenous networks of host cell can lead to cellular context dependent effects [7, 8]. For this reason, molecular parts and devices that are orthogonal to the cell native machineries with roles in either genetic transcription or protein translation have been created to enable predictable engineering of genetic circuits [9-13]. Demonstrations of layered genetic circuits in single cell, such as the execution of 4-input AND gate in bacteria [10] and biological half adder and half subtractor in mammalian cells [14] have revealed that orthogonal logic gates can be interlinked to perform digital operations of higher complexity and diversified outputs. While the capability to program cells with memory and decision making functions [15-19] presents many opportunities in biotechnological applications, a lack of formal understanding associated with genetic context dependent effects has limited progress in engineering biology. In this respect, two studies have shown that the 5’ untranslated region (5’-UTR) of mRNA can affect the temporal control of multigene operons or inverter-based genetic circuits, and RNA processing using CRISPR or ribozyme can serve as effective genetic insulators to buffer such context dependent effects [5, 20]. In this paper, we have sought to elucidate the limitations of engineering biology from an architecture point of view, with the aim of creating a set of engineering solutions for overcoming failure modes during the development of complex, synthetic genetic circuits.

### Design of Biological Half Adder

In this study we were interested in developing biological half adder in prokaryotic systems, particularly in microbes which exhibit much faster cell division and shorter cycle time – so that they can be broadly applied in different biotechnological applications. In contrast to the mammalian cell-based half adder, which is developed mainly for therapeutic and biosensing applications, a prokaryotic half adder can be used to enhance molecular process control and decision-making – for example in drug and biofuel production, biosensing, bioremediation [21], and probiotic engineering for the treatment of metabolic disorders [22], cancer [23] and infectious diseases [24, 25]. In digital processing half adders form the key building blocks for shift registers, binary counters and serial parallel data converters. Likewise in biological systems, a combination of half adders can be connected in various arrangements to regulate gene expression with diverse, digital-like performance. In doing so, biological systems can be made to interface with novel biomolecular devices, allowing the repurposing of cellular phenotype, as well as providing new platforms to probe and elucidate biological functions [26-28].

*Escherichia coli* was chosen as the designated chassis as it represents a model organism that can be easily manipulated - its inherent cellular processes are also well characterised. Fig. 1 shows the design of our biological half adder in a single prokaryotic cell. The half adder consists of 3 independent biologically-derived AND, OR and NOT logic gates - and a fourth AND logic function that is not a physical device, but a result of programmable decision making as a result of interconnecting logic functions (Fig. 1A). The σ^54^-dependent HrpRS regulation motif of *Pseudomonas syringae* T3SS secretion system was refactored for the design of the AND gate, as demonstrated in an earlier study [12]. The advantage is that the HrpRS AND gate offers dual layer of orthogonal control in *E.* coli host.

**Fig. 1.**
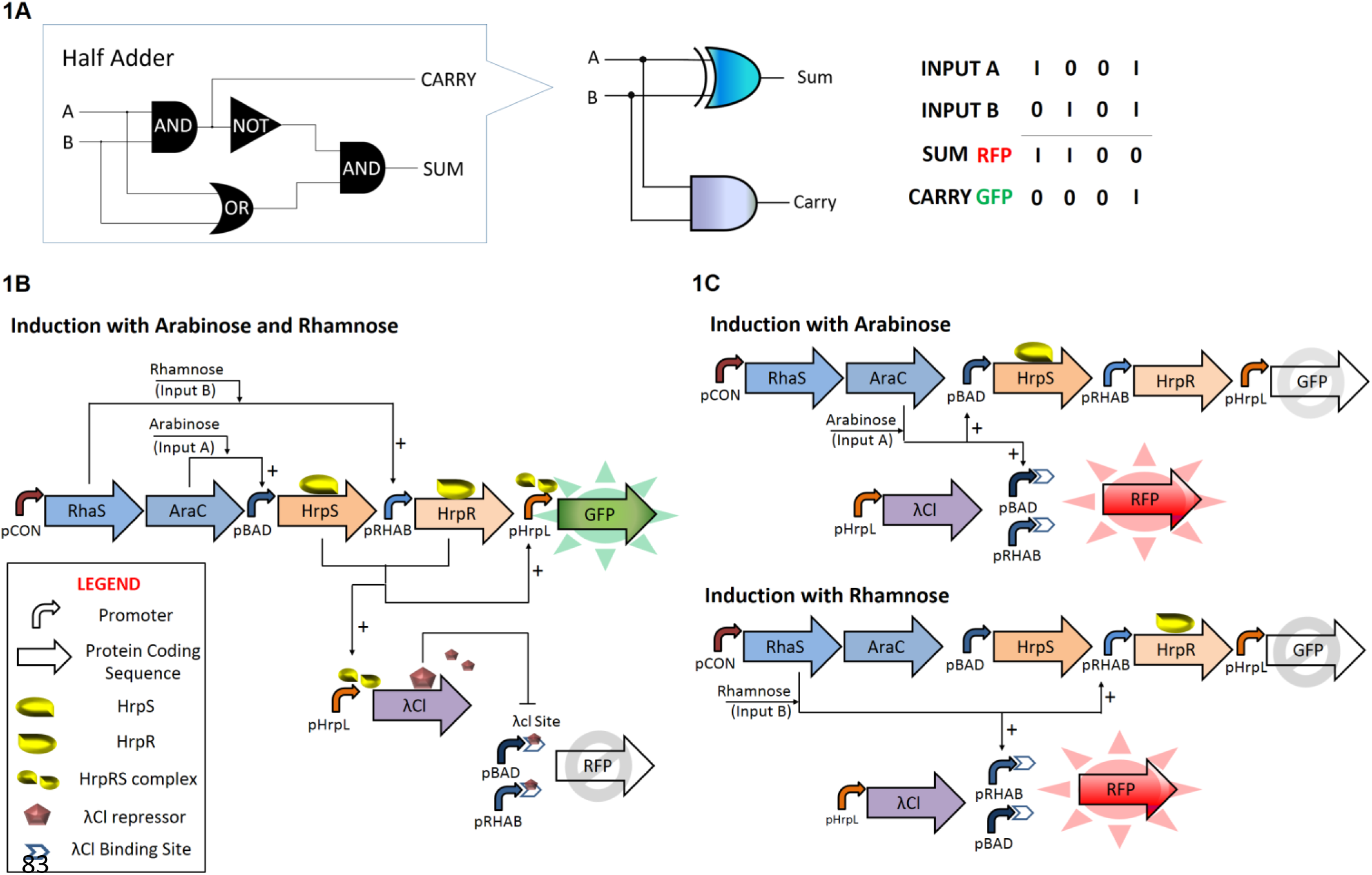
Simplified schematics of the biological half adder, comprising independent modules of the AND, OR and NOT gates layered in series and in parallel.

A. Logic output of biological half adder.
B. In the presence of two inputs, the AND gate is activated to produce GFP and lambda repressors, which further inactivates the OR gate to suppress RFP expression.
C. In the presence of either inputs singly, only the OR gate is activated to trigger RFP expression.

This means that (a) the majority of transcription events in *E. coli* occurs via σ^70^-dependent transcription, and (b) HrpRS transcription factors are absent in wild type *E. coli*. Transcription occurs when enhancer-binding proteins HrpS and HrpR, which are regulated by arabinose (input A) and rhamnose (input B) induction respectively, are coexpressed and bound to the upstream activator sites of pHrpL promoter. This binding event then triggers an ATPase-dependent conformational change within the promoter through a molecular interplay with the σ^54^-RNAP holoenyzme, thereby allowing RNA synthesis and elongation after the transcription start site. The OR gate generates mRNA transcript of the RFP gene upon induction with either arabinose or rhamnose. The NOT gate in the half adder design is a hybrid promoter consisting of λCl repressor binding sites downstream of the transcriptional start site (TSS) of OR logic gate. Unlike traditional NOT gates, which are designed to have transcriptional repressors competing for consensus RNAP binding sites, our NOT gate design functions as an orthogonal, molecular blocker to the RNA elongation process.

On induction with arabinose and rhamnose, the transcription factors AraC and RhaS, both of which are constitutively expressed in a single transcript by promoter pCON, associate with their corresponding inducers to activate expression of the enhancer-binding proteins HrpS and HrpR. This results in the activation of the AND logic and the concurrent synthesis of GFP reporter and lambda repressor (λCl) by the pHrpL promoter. Consequently, genetic events of the OR gate, which run in parallel with the HrpRS AND gate, is then turned off due to obstructive repression by λCl molecules. In all, the half adder demonstrates both AND (SUM Output) and XOR (CARRY Output) logic operations, the latter operation is a processed outcome achieved by sequential and parallel layering of AND, OR and NOT logic (Fig. 1B). By comparison, induction with either inducer singly will trigger only genetic operation of the OR gate, resulting in the synthesis of RFP reporter, but not GFP and λCl molecules (Fig. 1C). Finally, we also demonstrate the development of a single cell prokaryotic, half subtractor via slight modifications to the half adder circuit.

## MATERIAL AND METHODS

### Strains, plasmids and growth conditions

*E. coli* strain Top10 (Invitrogen) was used for all the cloning and characterisation experiments. The genes and oligonucleotides used in this study were synthesized by either Geneart (Life Technologies, Grand Island, NY) or Sigma Aldrich (St. Louis, MO). All the enzymes used in this study, including OneTaq and Phusion polymerase, T4 ligase, EcoRI, XbaI, SpeI, PstI and DpnI, were obtained from New England Biolabs. Chloramphenicol (35μgml^-1^) and ampicillin (100 μgml^-1^) were added to culture media for experiments involving pSB1C3 and pSB4A5 plasmid vectors, where appropriate. In all the characterisation experiments, cells were inoculated from freshly transformed plates were grown in 2ml LB (Miller, BD Bioscience, San Jose, CA) with appropriate antibiotic in 50ml Falcon tubes overnight at 37oC with 225rpm shaking unless otherwise stated. Overnight cultures were then diluted to OD_600_ ∼ 0.002 in 5ml LB antibiotic and further grown to a final OD of 0.5± 0.05 with the same culture conditions (37°C and 225rpm shaking). Harvested cells were kept on ice until induction. All inducers used in this study were purchased from Sigma Aldrich with final concentration ranging from 0 to 28mM.

### System assembly

All genes from *E. coli* (*pBAD*, *pRHAB*, *araC* and *rhaS*) were cloned from genomic DNA of strain MG1655 (ATCC 700926). *hrpS*, *hrpR* and *pHrpL* were cloned from an earlier study [12] while *pCON* (Bba_J23101), double terminator (Bba_B0015), GFPmut3b (Bba_E0040), RFP (Bba_E1010) and *λCl* (Bba_C0051) were cloned from the Biobrick registry. PCR was performed using Phusion DNA polymerase in a dual cycle PCR programme at annealing temperatures of 53°C and 60°C for the first 7 and subsequent 20 cycles, respectively. Biological parts were spliced by overlap extension PCR and ligated to vectors pSB4A5 (low copy, pSC101 replication origin) and pSB1C3 (high copy, pMB1 replication origin) using XbaI and PstI restriction sites. Composite systems with two or more biological modules were sequentially assembled as previously described [25].

### Parts mutation of λCl repressor binding sites

To obtain sequence variants of λCl repressor binding sites, PCR with randomised primers and Phusion DNA polymerase were performed on pHrpL-λCl-pBAD-Cl2A template with primers 5’ - ttc<u>gaattcgcggccgcttctagag</u>gccggattat and 5’-gct<u>actagt</u>atatNNNNNNNNccggtgatatatggagaaacagta (restriction sites underlined). The resultant amplificons (∼1.4kb) were then ligated upstream to pSB1C3 vector containing an RFP reporter and transformed to competent cells carrying HrpRS AND gate modules in pSB4A5. Single colonies of uniform size were inoculated into 96 well microplate loaded with 200ul LBAC (LB with chloramphenicol and ampicillin) and grown in microplate incubator set at 37°C with 750rpm shaking for 6 hours. Accordingly, cultures in each well were triplicated and diluted 10x into 200ul LBAC with 3.5mM arabinose and 28mM rhamnose, 3.5mM arabinose, and no arabinose in the same growth condition. Evolved mutants were identified by observable differences in RFP expression and inhibition after 6 hours of induction using Fluostar OPTIMA microplate reader (BMG Labtechnologies). Validation and characterisation of isolated candidate parts was independently performed in 175ul LBAC in 1.5ml microcentrifuge tubes after 4 hours induction at 37°C and 1000rpm shaking under four different logic conditions.

### Modelling of AND, OR and NIMPLY logic gates

To enable model-driven design synthetic biological systems, we examine the effect of ribosome binding sites (RBS) on the steady state transfer function of input switch devices. By analysing reference data [12] that had previously characterise the input-output relationship of genetic switches in the form of Eqn. 1, we observed that parameters that are most sensitive to changes in RBS are parameters A and B. Hence, by knowing the relative output of switch devices with weaker RBS by either prediction from reliable software or by single experimental measurement of device’s output at input maximal, the parameters A and B can be scaled proportionally to obtain *a priori* parameters that accurately predict the transfer function of other devices with weaker RBS (Supplementary Fig. 1A). We validated our approach with previously published data sets (Supplementary Table 1) and showed that the transfer function of input devices pLuxR (Supplementary Fig. 1B) and pBAD (Supplementary Fig. 1C) with different RBS can be reliably estimated without additional experimentation. MatLab modelling scripts are available in Supplementary Material II.

**Table 1.** Failure modes and engineering solutions for the design and built of layered genetic circuits in single (bacterial) cell.

| Device | Failure Mode | Engineering Solution | Fig / [Ref] |
| --- | --- | --- | --- |
| Input switches | **Genetic crosstalk:** Input switch devices cross-talk with one another. | Check pairwise compatibility by placing GFP and RFP under the regulation of each input switch device<br>Perform mutagenesis on promoter or DNA-binding protein to identify orthogonal pairs. | S4<br>Ref [28, 29, 31, 50] |
| **AND gate** | **Stoichiometric mismatch:** Amount of AND gate's transcription activators are disproportionately matched, resulting in "leaky" AND gate.<br>**DNA supercoiling:** $\sigma 54$ AND gate promoter is turned on by the DNA supercoil effects of upstream $\sigma 54$ promoter | Characterise the expression profile of input genetic switches with different RBS and input the resultant transfer function equations into a steady state AND gate computational model. Match AND gate sub-modules to obtain stoichiometric balance using this forward engineering approach.<br>Insulate $\sigma 54$ promoters using different plasmid vectors. | S11<br>S5 |
| **OR gate** | **Stoichiometric mismatch:** Outputs from input device I and II are disproportionately matched, resulting in skewed OR gate.<br>**Transcription interference:** Tandem promoter OR gate design fails due to downstream DNA sequence acting as a repressor to upstream promoter. | Characterise the expression profile of input genetic switches with different RBS and input the resultant transfer function equations into a steady state OR gate computational model. Match OR gate sub-modules to obtain stoichiometric balance using this forward engineering approach.<br>Characterise different permutation of tandem promoter OR gate to identify the optimal genetic architecture.<br>Separate OR gate promoters into distinct expression cassettes. | S12,S13<br>3A,3C<br>3A,3C |
| **Layering OR-NOT into NIMPLY gate** | **Insufficient repression:** Placing single repressor binding site downstream of inducible promoter cannot fully repress gene expression.<br>**Translation interference:** Placing repressor binding sites downstream of inducible promoter creates extensive 5'UTR structural effects. | Increase repression efficiency by introducing additional repressor binding sites to the NOT gate. Note that the introduction of extra repressor binding sites may also lead to extensive 5'UTR effects.<br>Attenuate expression "leakiness" by using weaker RBS for the NOT gate<br>Perform mutagenesis to relieve RNA hairpin structures at selected sites.<br>Use RNA processing tools to remove undesired 5'UTR sequences. | 4A,4C<br>4B<br>S6<br>Ref [5, 20] |
| **Layering AND-OR-NOT into XOR gate** | **Insufficient repression:** Insufficient transcription repressors are generated by upstream genetic circuit to stop transcription elongation, level mismatch.<br>**Translation interference:** Placing repressor binding sites downstream of OR gate tandem promoter creates extensive 5'UTR structural effects. | Reduce repressors required in NOT gate by designing repressor binding sites such that it is immediately downstream of transcription start site.<br>Increase production of repressor in the AND gate by expressing transcription repressors in high copy plasmid.<br>Separate OR gate promoters into distinct expression cassettes.<br>Use RNA processing tools to remove undesired 5'UTR sequences. | 5<br>2D,5<br>5<br>Ref [5, 20] |

The transfer functions of input switch devices used in this work with strong RBS were empirically fitted into the Hill-like equation (Eqn. 1), while those of input switch devices with weak RBS were predicted using the validated method as discussed above. Supplementary Table 3 shows the empirical transfer function parameters of the various input switch devices. AND and OR gate profiles were then modelled and predicted using these parameters and equations as shown in Supplementary Materials Eqn. 7 and 10. Supplementary Fig. 11, 12 and 13 show the predicted normalised output of the HrpRS AND gate and various OR gates combination. NIMPLY gate was empirically modelled using Supplementary Materials Eqn. 2 and parameters from Supplementary Table 2. Supplementary Fig. 10 shows the predicted output of NIMPLY gate.

### Characterisation and orthgonality testing of input switch devices

To characterise input switch devices, reinoculated cultures at OD_600_ ∼0.5 were transferred to black, flat-bottom 96-well plates (Greiner Bio-One, cat. no. 655090) in aliquots of 150μl for induction with rhamnose or arabinose in serial 2 fold dilution with highest inducer concentration at ∼28mM (i.e. 0.00M, 8.38E-07M, 1.68E-06M, 3.35E-06M, 6.70E-06M, 1.34E-05M, 2.68E-05M, 5.36E-05M, 1.07E-04M, 2.14E-04M, 4.29E-04M, 8.58E-04M, 1.72E-03M, 3.43E-03M, 6.86E-03M 1.37E-02M, 2.75E-02M). Plates were then sealed with gas-permeable foils and incubated at 37°C with 750rpm shaking for 3 hours. Fluorescence and optical density data were collected using Fluostar Optima microplate reader (BMG Labtech.) and zeroed with blank LB media with antibiotic to remove background fluorescence and OD_600_. All results were normalized with OD_600_-estimated cell density (validated with viable cell counts) and provided in arbitrary units. In the orthogonal testing of input switch devices, the above procedures were repeated with constructs that contain pRHAB-RFP-pBAD-GFP, with fixed concentration of 0.02% arabinose or rhamnose added as appropriate. The experimental results were fitted using an empirical mathematical model [25] (Hill equation),

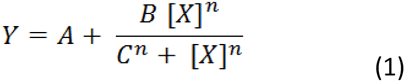

Equation 1 models reporter output (*Y*) as a function of input concentration of inducer ([X]). The four parameters (*A, B, C, n*) were estimated to obtain the best fit curve by performing a nonlinear curve fitting using the experimental results. This curve fitting was performed using the nonlinear least square fitting functions in MATLAB Curve Fitting Toolbox (The Mathworks, Natwick, MA, USA).

### Characterisation of AND, OR, NIMPLY, XOR, half adder and half subtractor

To characterise the steady state profile of AND and OR logic devices, reinoculated cultures at OD_600_∼0.5 were transferred to black, flat-bottom 96 well plates (Greiner Bio-One, cat. no. 655090) in aliquots of 150μl for induction with varying concentrations of rhamnose and arabinose ranging from 8.38E-07M to 2.74E-02M. The plates were sealed with gas-permeable foils and incubated at 37°C with 750rpm shaking for 3 hours. Fluorescence and optical density data were collected using Fluostar Optima microplate reader (BMG Labtech.) and zeroed with blank LB media.

To characterise the steady state profile of NIMPLY, XOR, half adder and half subtractor logic devices, the above procedures were repeated with slight modifications to reduce evaporation losses in constructs with weaker RBS-RFP modules. Briefly, reinoculated cultures were dispensed in 175μl aliquots into 1.5ml capped-tubes and induced with varying concentrations of rhamnose and arabinose, as above. The aliquots were incubated on a thermomixer platform (Eppendorf) set at 37°C with 1000rpm shaking for 4 hours. 150μl aliquots from each tube were transferred to black, flat-bottom 96 well plates and assayed for fluorescence and optical density with Synergy HT or H1m microplate readers (Biotek Instruments Inc.). To assess the digital performance of all logic devices, cell cultures were separately induced with water, 28mM rhamnose or/and 7mM arabinose in four different logic conditions. The induced cultures were incubated in the respective conditions as described above and assayed for fluorescence. All results were normalized with OD_600_-estimated cell density and provided in arbitrary units.

### Fluorescence imaging of AND & OR gates

For the acquisition of fluorescent images in AND and OR logic devices, reinoculated cultures were transferred to 50ml tubes in aliquots of 5ml and separately induced with water, 28mM rhamnose or/and 7mM arabinose in four different logic conditions overnight. After 15 hours, cell pellet were harvested and transferred to 1.5ml tubes for fluorescent imaging with suitable filters. Images were acquired with high mega-pixel mobile phone camera.

### Flow cytometry

Reinoculated cultures were dispensed in 175μl aliquots into 1.5ml capped-tubes and separately induced with water, 28mM rhamnose or/and 7mM arabinose in four different logic conditions. The aliquots were grown on a thermomixer platform (Eppendorf, Germany) set at 37°C with 1000rpm shaking for 4 hours. Before assay, 5μl culture from each sample were diluted 200x in 0.22μm filtered DI water (pH 7). All expression data were collected using BD LSRFortessa X-20 flow cytometer (BD Biosciences, San Jose, CA) with a 488nm argon excitation laser, and 530nm±30 (FITC) and 610nm±20 (PE-CF594) emission filters. The data were gated using both forward (550v, threshold 1500v) and side scatter (310v) with the neutral density filter removed. At least 10,000 events were recorded per sample. FITC and PE594 channels were set at 466v and 852v respectively. Data analysis was performed with FlowJo (TreeStar Inc., Ashland, OR).

## RESULTS

### Characterisation of Input Devices

The choice of input signals presents the first possible complication in terms of parts modularity. For this reason, genetic circuits of higher complexity with multiple inputs often utilise promoter systems which are activated by inducers of vastly dissimilar chemical nature, namely IPTG, tetracycline, arabinose, 3OC12HSL and C4HSL. Previous studies have shown that a subset of quorum sensing promoters can be activated by homoserine lactone inducers of similar carbon chain length [29, 30]. Likewise, wild type pBAD promoter is affected by lactose analogues, requiring further mutagenesis to avoid crosstalk inhibition [31]. Instances of cross-phosphorylation have also been observed in two component signal transduction systems between otherwise distinct pathways [32]. Thus, it is important that inducible input devices are carefully characterised for their steady state transfer function and pairing compatibility before further assembly into higher ordered logic devices.

While previous studies with pRHAB promoter involved genetic circuits that include both RhaR and RhaS transcription factors [33-35], in this paper we demonstrate that the rhamnose inducible promoter pRHAB requires only RhaS for full activation and displays tight regulation even when RhaS is overexpressed. Supplementary Fig. 2C and S3C show the steady state transfer functions of input device A, pBAD (Supplementary Fig. 2A) and input device B, pRHAB (Supplementary Fig. 3A) expressing RFP under strong RBS by their corresponding inducers, respectively.

**Fig. 2.**
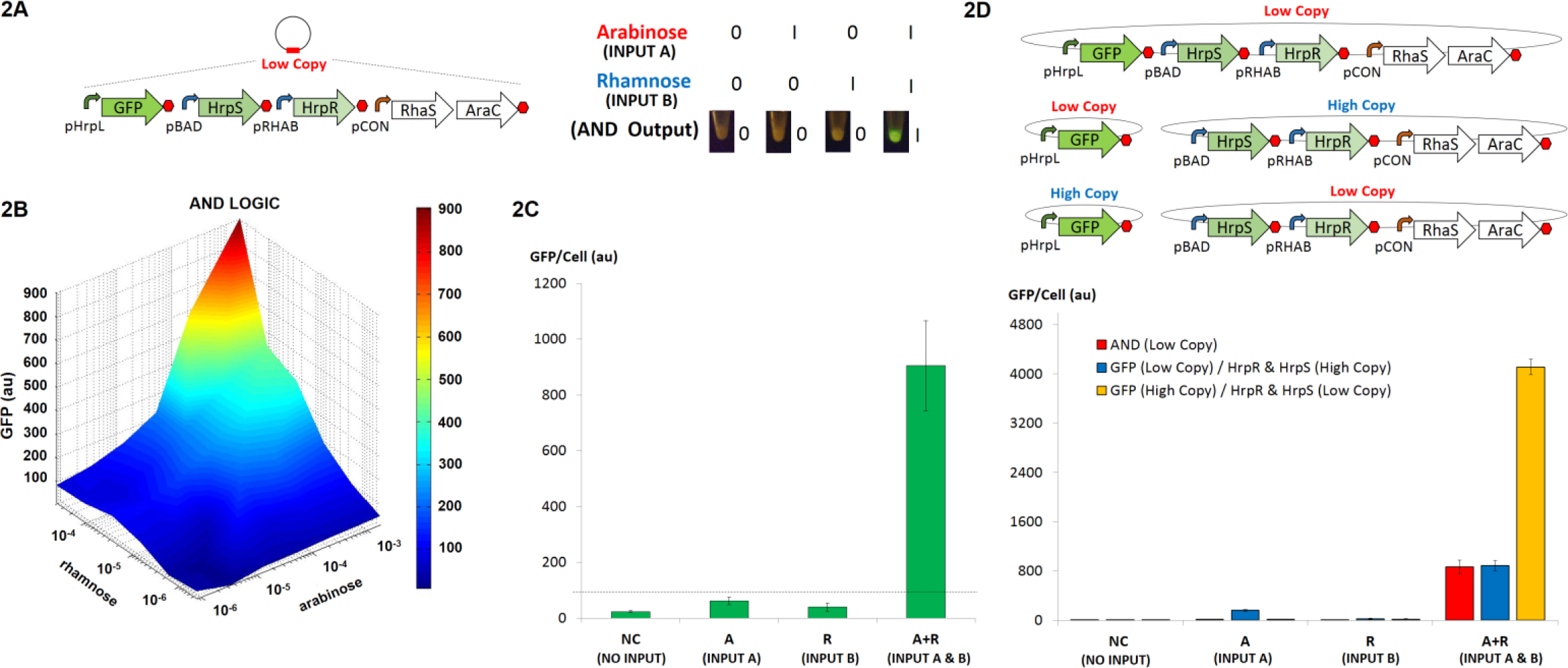
Design and characterization of the biological AND gate.

A. Design and logic output of Hrp-based AND gate. The AND gate comprises of HrpS and HrpR transcription factors that are unregulated under the control of pBAD and pRHAB promoters, respectively. In the presence of both inputs HrpRS jointly bind and induce conformational change in the pHrpL promoter, thereby enabling DNA transcription and the expression of GFP reporter.
B. Steady state profile of the AND gate for various concentrations of arabinose (input A) and rhamnose (input B).
C. Digital performance of AND gate at steady state.
D. Characterization of the Hrp-based AND gate in both high and low copy plasmids. The input devices generating HrpRS transcription factors and pHrL-GFP reporter module are placed in plasmids of different copy numbers to study the effect of plasmid copy on precision control and tuning of Hrp-based AND gate. Error bars represent the standard deviation of three independent experiments.

**Fig. 3.**
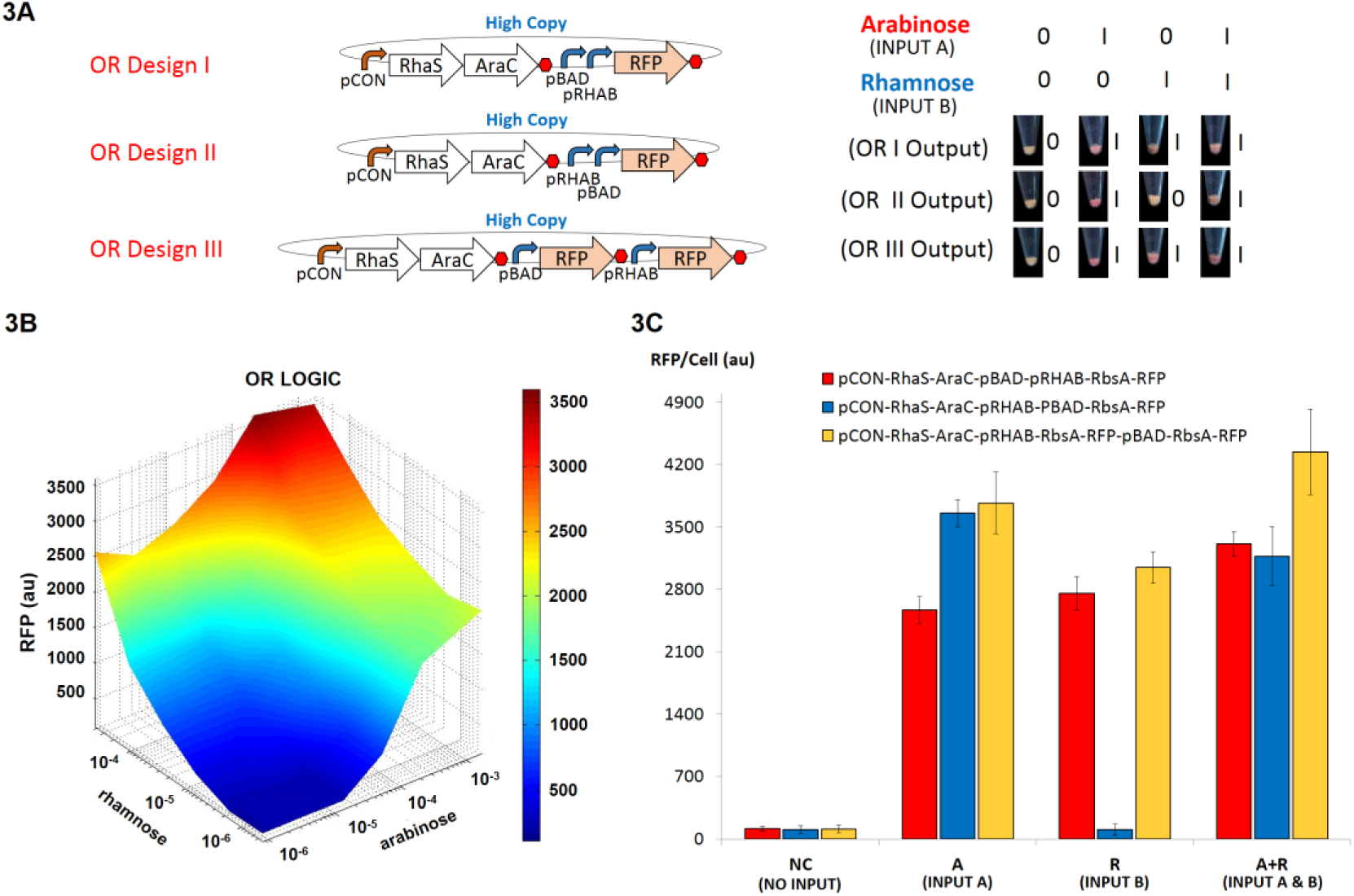
Design and characterization of biological OR gates.

A. The genetic blueprint and logic output of three OR gate designs. Designs I and II are tandem promoters in opposite arrangement, while design III expresses RFP reporter in two distinct transcripts. Only design I and III are functional OR gates that generates RFP in the presence of either inputs.
B. Steady state profile of OR gate I for various concentrations of arabinose (input A) and rhamnose (input B).
C. Digital performance of OR gates at steady state. Error bars represent the standard deviation of three independent experiments.

To examine the possibility of genetic cross-communication, we constructed genetic circuits that couple GFP production to pBAD activation and RFP production to pRHAB activation. The results show that varying concentration of arabinose did not activate pRHAB promoter activity (Supplementary Fig. 4A). A similar trend was observed in pBAD promoter with rhamnose (Supplementary Fig. 4B). Interestingly, the simultaneous introduction of both sugars modified the transfer function of each promoter slightly, which may be a result of differential cell growth, sugar import rate or antagonistic effect of one sugar to another. This effect, however, is insignificant as definite ON and OFF switch behaviours are apparent - thereby confirming the pairing compatibility of pBAD and pRHAB promoters.

**Fig. 4.**
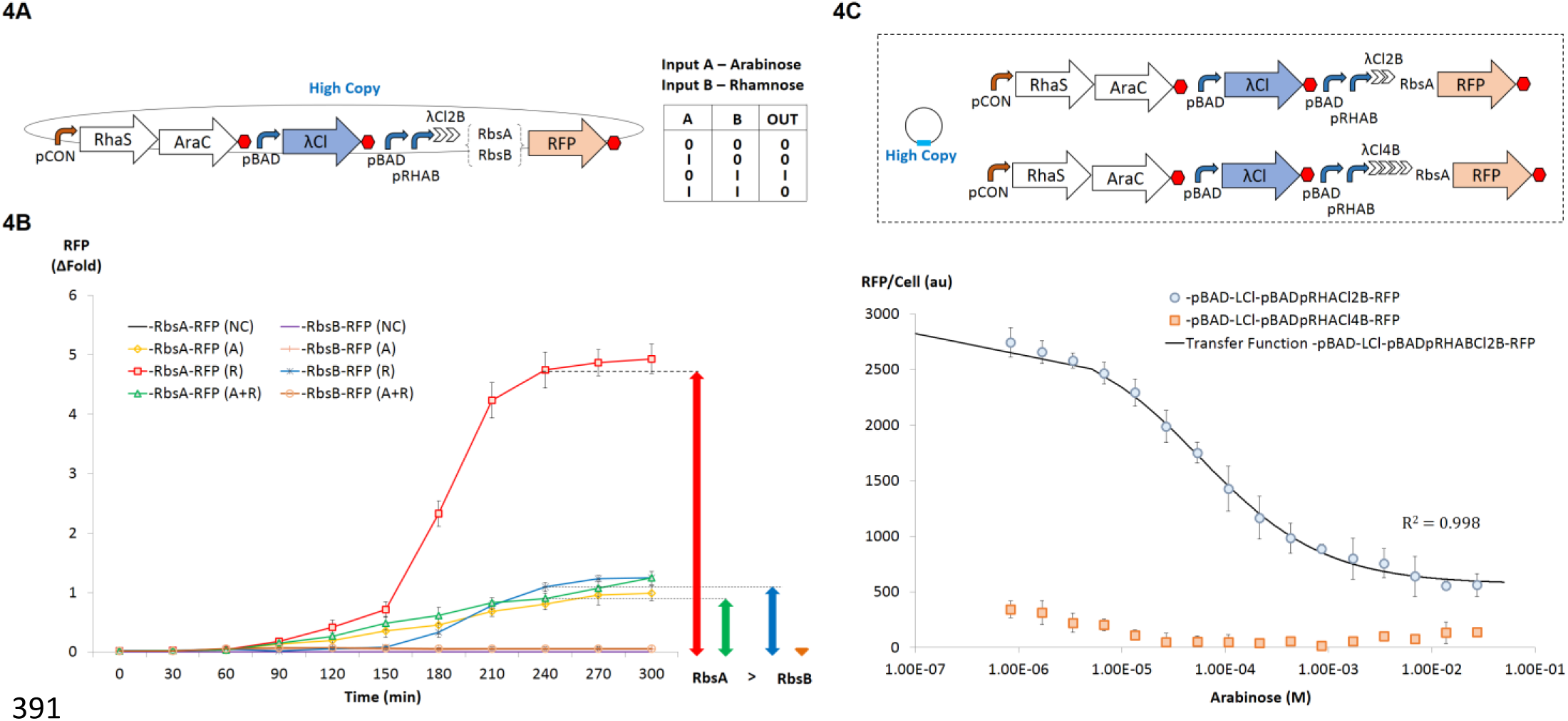
Design and characterization of the biological NIMPLY gate.

A. Genetic blueprint and logic output of NIMPLY gate. The NIMPLY gate is designed by incorporating synthetic lambda repressor binding sites downstream of OR gate promoters and regulating the expression of lambda repressors through the pBAD promoter. RFP is expressed only in the presence of input B, rhamnose.
B. Characterization of NIMPLY gate with different ribosome binding sites. At steady state NIMPLY gate which utilizes a weaker ribosome binding site (RbsB) directly upstream of the RFP reporter (denoted by black stars, orange circles and purple crosses) exhibits better control and reduced expression leak, as compared to the NIMPLY gate design that contains a stronger ribosome binding site (denoted by red squares, green triangles and blue diamonds). Expression leakiness in circuits with strong and weak ribosome binding sites after 4 hours are denoted by green and orange arrowheads, respectively. Constructs that were singly induced with input B, induced with both inputs A and B, and uninduced are represented by R, A+R and NC as shown.
C. Characterization of NIMPLY gates with two (blue circles) and four (red squares) lambda repressor binding sites. The black line represents empirically-derived transfer function for the construct with dual lambda repressor binding sites, as described by the equation provided. Constructs were induced with a fixed amount of rhamnose (input B) and titrated with various concentrations of arabinose (input A). An increased number of repressor binding sites disrupted the NIMPLY gate, possibly due to pronounced effect of 5’ mRNA secondary structures. Error bars represent standard deviation of three independent experiments.

### Design and Characterisation of AND Logic Gate

Designs of highly modularised, prokaryotic AND logic devices have hitherto involved the use of multiple plasmids [10, 12, 16, 36, 37]. In this work, we assembled AND logic gate in a single plasmid. This procedure has enabled us to localise AND logic gate in a single vector, and facilitated the downstream troubleshooting and tuning of layered genetic circuits.

To develop the AND logic component of the half adder, we systematically designed and assembled refactored modules of the HrpRS transcription machinery into a low copy plasmid (Fig. S2A). The module which expressed GFP from pHrpL promoter was assembled upstream of pBAD-HrpS and pRHAB-HrpR modules to attenuate genetic context dependent effects that might arise from transcriptional overrun of the stronger pBAD and pRHAB input expression modules as a result of inefficient transcription termination. While designing the GFP producing module in a bidirectional permutation is usually a better solution, this option was not tested in our study as the downstream pBAD promoter is a weak constitutive promoter in the reverse complement direction. Thus, placing the pHrpL-GFP module before pBAD in either the reverse or reverse complement arrangement may result in antisense-GFP interference or the occurrence of leaky AND gate. The steady state profile of the functional AND gate was characterised by titrating with a varying concentration of arabinose (input and rhamnose (input B) as shown in Fig. 2B. Results of the engineered AND gate correlated well with our steady state computational model (Supplementary Fig. 11), which was applied to match biological modules making up the AND gate. Likewise, the “on” and “ off” digital performance of the AND gate at steady state was qualitatively and quantitatively assessed by introducing inputs well above switch points under four different logic conditions (Fig. 2A and 2C). The results show that the AND gate was only activated in the presence of both inputs with >800au (relative fluorescence unit) expression increase, as compared to the condition where only single input is present (or no inputs).

To assess the effect of plasmid copy number on the performance of the AND gate, modules were constructed which generate the HrpRS transcription activators (pBAD-HrpS-pRHAB-HrpR). This produces a GFP output (pHrpL-GFP) into separate low and high copy plasmids (co-transformed the plasmids into E. coli cells).The relative GFP output of each system was measured (Fig. 2D). The results show that the AND gate system with the GFP-producing module in high copy plasmid and HrpRS transcription activators in low copy plasmid produced a >4 fold greater GFP output than AND gate systems with GFP-producing module in low copy plasmid and HrpRS (as compared to transcription activators in either low or high copy plasmids). The result indicates that a higher concentration of HrpRS transcription activators, above the saturation limit of the pHrpL promoter, do not produce a greater GFP output. It is likely that the transcriptional output of the HrpRS AND gate is limited by the strength of the weak pHrpL promoter. Hence, the conclusion is that when pHrpL-GFP module was expressed in high copy plasmids, intracellular availability of pHrpL promoters were increased - resulting in the amplification of GFP output.

### Design and Characterisation of OR Logic Gate

Genetic OR gates can be achieved by designing tandem promoter genetic circuits or by expressing target gene in two discrete expression cassettes. Nonetheless, tandem promoter OR gate circuits may fail when repression of downstream promoter prevents the proper functioning of the upstream promoter [38]. To develop the OR logic gate of the half adder, three prototype designs were constructed; two of which comprised of pBAD and pRHAB promoters in different tandem arrangements upstream of an RFP reporter gene with strong RBS, and a third design that produces RFP in two distinct expression cassettes (Fig. 3A). The three OR gate designs were then introduced with input A and B above their switch points and assessed for the respective RFP outputs (Fig. 3A and 3C). The results show that designs I and III are functional OR gates with >2500au higher RFP expression when either or both inputs are present. In our computational model, the total amount of RFP expression was approximated by the sum of RFP amounts produced from individual pBAD and pRHAB promoters. Although the model predicts well from low to medium range induction levels, our assumption was not valid at very high induction levels, in which lesser RFP expression was observed than predicted. It is possible that at very high induction level, the transcription and translation machinery in cells are fully saturated, thereby imposing metabolic burden on the cells and limiting protein production [39]. The OR gate design II, which composed of pRHAB promoter upstream of pBAD promoter and RFP reporter was activated only in the presence of rhamnose, but not arabinose. Our results agree with previous finding that no expression was detected when pBAD promoter was fused downstream of tetracycline-inducible pTET promoter and upstream of a YFP reporter [38]. The conclusion is that it is likely that this observation is an effect of the AraC transcription factor - which can function as both repressor and activator. In the absence of arabinose, AraC when over expressed, remains bound to operator sites that induce DNA looping of the pBAD promoter, thereby obstructing the elongation of mRNA by initiated RNA polymerase. As will be shown in the next section, in order to layer OR gate design I into other logic devices, the construct was characterised for its steady state profile by titrating with varying concentration of arabinose and rhamnose (Fig. 3B). Results of the engineered OR gate generally correlated well with our steady state computational model (Supplementary Fig. 12), which was applied to match biological modules making up the OR gate.

### Genetic Context Effect of σ54-dependent pHrpL promoter

To enable sufficient expression of the λCl repressor by an AND gate system, the gene encoding for λCl repressor was assembled downstream of σ54-dependent pHrpL promoter on a high copy plasmid. Fortuitously, we discovered that pHrpL promoter located downstream of another pHrpL expression cassette can be turned on even in the absence of its cognate HrpRS transcription factors (Supplementary Fig. 5C). The converse is not true for an upstream pHrpL promoter (Supplementary Fig. 5B). Negative controls with just the GFP reporter or RBS-λCl gene upstream of pHrpL-GFP module confirmed that pHrpL promoter alone is not leaky and that cryptic promoter is absent in the λCl gene (Supplementary Fig. 5A and 5D). To buffer against this genetic context dependent effect of the pHrpL promoters, pHrpL-GFP and pHrpL-λCl modules were assembled on separate plasmids. This successfully prevented the genetic interference of both pHrpL expression modules on each other (Supplementary Fig. 5E and 5F). Supplementary Fig. 5G shows a quantitative assessment of pHrpL promoter activation due to the presence of another upstream pHrpL promoter and the use of plasmids as genetic insulators.

**Fig. 5.**
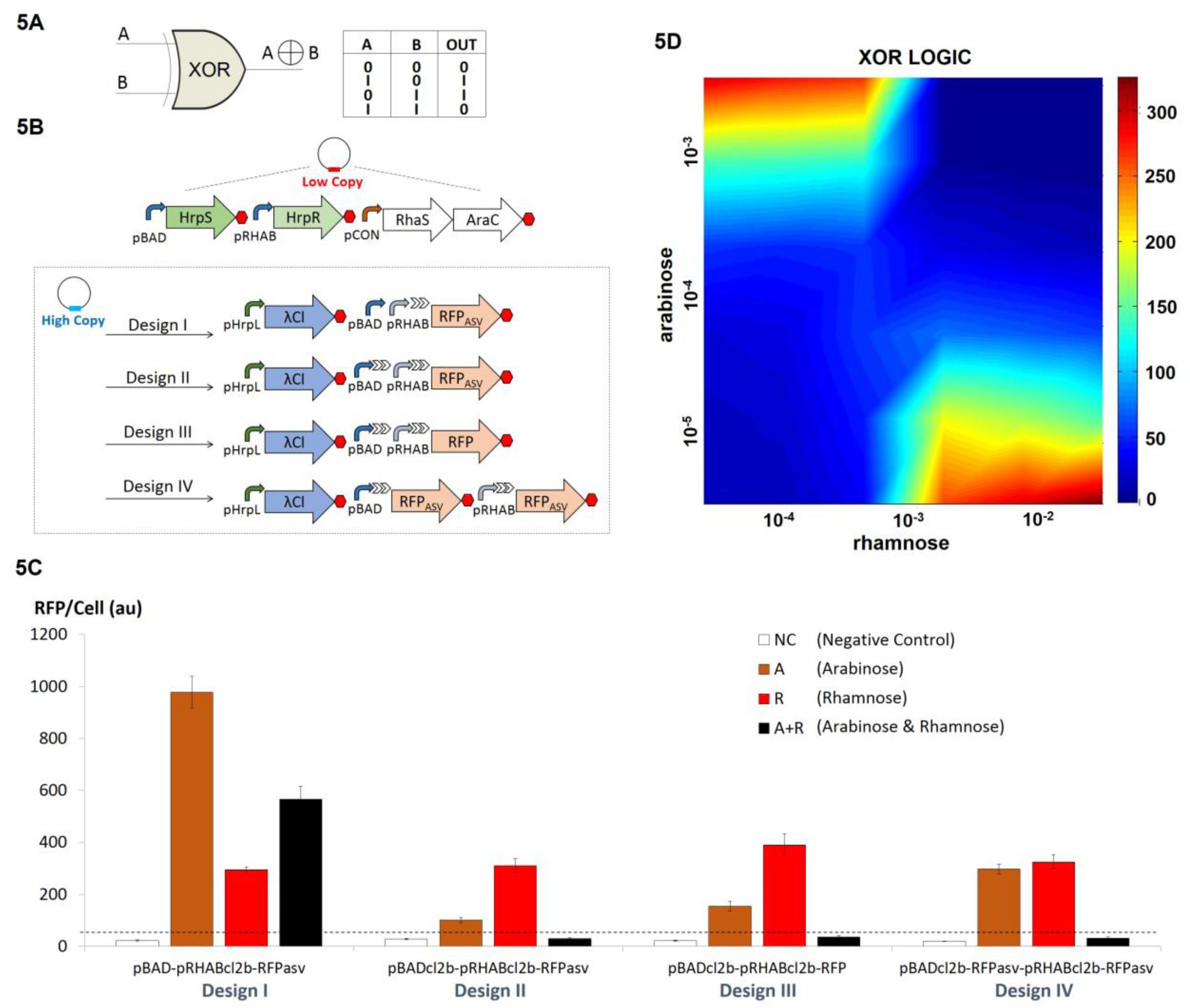
Design and characterization of biological XOR gates.

A. The logic output of XOR gate.
B. Genetic blueprint of four biological XOR gate designs. The XOR gate comprises of serially layered AND, NOT and OR gates. HrpRS transcription factors are carried in a low copy plasmid, while pHrpL-λCl and distinct modules of OR gates with lambda repressor binding sites expressing RFP reporter are carried in high copy plasmids. Design I comprises tandem promoters with repressor binding sites downstream of pRHAB promoter and a RFP reporter engineered with the ASV protein degradation tag. Designs II and III comprise tandem promoters with repressor binding sites downstream of each promoter and RFP with and without the ASV degradation tag respectively. Design IV is modified from design II with RFP expressed in two disparate transcripts.
C. Digital performance of various designs of biological XOR gates at steady state.
D. The steady state profile of XOR gate IV for various concentrations of arabinose (input A) and rhamnose (input B). Error bars represent the standard deviation of four independent experiments.

### Design and Characterisation of NOT and NIMPLY Logic Gates

As part of the development of XOR logic operations of the half adder, repressor binding sites are required downstream of the OR gate promoters. To examine the minimal number of λCl repressor binding sites required for effective repression, single λCl operator site and dual λCl operator sites of perfect dyad symmetry were fused downstream of pBAD promoter, before the RFP gene [40]. The repressibility of both circuits was tested by generating λCl repressors from HrpRS AND gate in a separate plasmid. Negligible repression was observed when only one λCl repressor operator site was present. In the presence of two operator sites of perfect dyad symmetry, RFP expression from pBAD promoter was greatly attenuated - even when λCl repressor was not synthesized. We postulate that the observed reduction of RFP expression might be caused by the presence of secondary hairpin structures immediately downstream of TSS acting as *pseudo* transcription terminator or locking RBS in conformations that prevented translation initiation (Supplementary Fig. 6A).

**Fig. 6.**
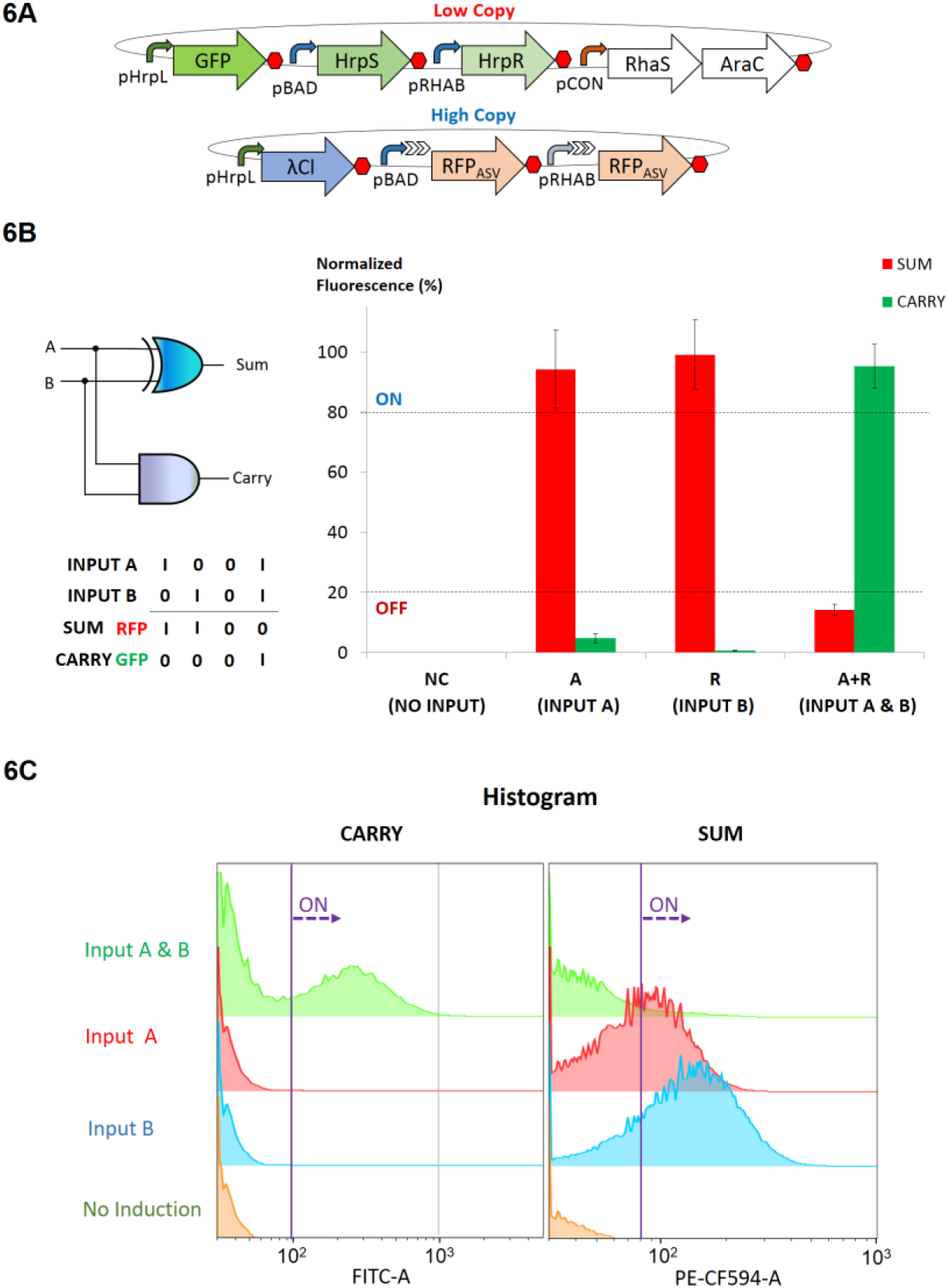
Design and characterization of the biological half adder.

A. Genetic blueprint of the half adder.
B. Digital performance of the half adder at steady state.
C. Flow cytometry analysis of the half adder. The Y axis coordinate represents population count, while FITC-A and PE-CF594-A represent channels that detects GFP and RFP fluorescence respectively. Population shifts to the right represent ON behaviour. Error bars represent the standard deviation of four independent experiments.

In order to examine this further, random mutagenesis on the natural sequence of the λCl repressor operator sites were performed and screened for mutants with significant difference in RFP expression levels, in the absence and presence of λCl repressor. Accordingly, an evolved candidate (Cl2B) with 4 mutations in the inverted sequence of the λCl repressor binding (Supplementary Fig. 6B) was obtained. Sequence comparison with the original λCl repressor binding sites (Cl2A) with the evolved candidate revealed that the directed evolution process had eliminated the effect of secondary hairpin structures from 7 to 3. Next, the efficiency of λCl-mediated transcription termination in the context of a genetic NIMPLY gate was studied. This was achieved by placing repressor binding sites directly downstream of tandem pBAD-pRHAB promoters and generating λCl repressors from a separate pBAD expression cassette.

Two NIMPLY logic circuits were developed which generated RFP transcripts with strong and weak RBS. Both NIMPLY logic circuits were then tested in the presence and absence of input A (arabinose) over time with input B (rhamnose), both above switch point (Fig. 4A). Temporal analysis of the NIMPLY logic circuits showed that there was no significant delay in layering NOT gate downstream of an OR gate (Fig. 4B). However, apparent delay in total amount of mature RFP was observed when a weaker RBS was used to initiate the translation of RFP gene. The results also showed that while NIMPLY logic can be achieved from both circuits, the system with the strong RBS exhibited a higher order of expression and leakiness as compared to that which translated RFP from weaker RBS. This leads to the conclusion that the choice of a particular RBS can be used as a signal moderation technique in order to achieve a balance between precision tuning and output gain in layered logic gates. In an attempt to alleviate expression leakiness from the NIMPLY gate with strong RBS, an additional pair of λCl repressor binding sites with imperfect dyad symmetry were introduced downstream of pBAD-pRHAB-Cl2B, and before the RBS-RFP module. However, the presence of 4 λCl binding sites completely inhibited RFP expression, resulting in the failure of the NIMPLY gate (Fig. 4C). It is likely that this failure could be an effect of pronounced 5’ UTR secondary structures formed due to the repeated use of identical λCl repressor binding sites.

### Design and Characterisation of XOR Logic Gate

In order to develop the XOR component of the half adder, we assimilated and tested a combination of AND, OR, and NOT logic gates in four different genetic circuits. In all the designs HrpRS transcription activators were expressed from low copy plasmid to drive the synthesis of λCl repressors from pHrpL promoter in high copy plasmids (Fig. 5B). OR and NOT biological modules were assembled in the same high copy plasmid downstream of pHrpL-λCl module. In design I, an OR gate comprising a tandem arrangement of pBAD, pRHAB and λCl repressor binding sites was used to express *ssrA*-tagged, short-lived RFP (RFP_asv_) - one of the most well characterised protein degradation system in *E.coli* [41]. In design II we created hybrid promoters of pBAD and pRHAB by incorporating λCl binding sites downstream of both promoters before connecting them in tandem to elicit hypothetical OR logic as similar to design I. Design III was modified from design II to express long-lived RFP. To overcome possible complications from 5’ UTR secondary structures - due to presence of multiple λCl binding sites within the same mRNA transcript, design IV, which comprised of synthetic hybrid promoters of pBAD-Cl2B and pRHAB-Cl2B expressing RFP_asv_ in two discrete expression cassettes was also developed.

Accordingly, only design IV was able to achieve well-balanced outputs which accurately described XOR logic operations (Fig. 5C). While design I demonstrated the strong suppression of RFP output in the presence of both inputs (arabinose and rhamnose), when characterised as an NIMPLY gate (as described earlier), the same design failed to function in the context of XOR gate in which a weaker pHrpL promoter was used to drive the synthesis λCl repressors instead of the strong pBAD promoter. Interestingly, the results imply that when employing transcription repressors as molecular blockers to mRNA elongation, a higher concentration of λCl molecules is needed to completely suppress transcription as λCl binding sites are engineered further away from the transcription start site. This observation may be an effect of RNAP gaining momentum as it runs down template DNA to perform transcription, inadvertently enabling RNAP to continue its course of action as a result of the inadequacy of “molecular brakes”.

While designs II and III, that were developed with λCl binding sites downstream of both pBAD and pRHAB promoters, exhibited a slight semblance of XOR logic operations, the presence of multiple, repeated sequences of λCl binding sites in the transcript generated from the pBAD promoter greatly reduced the RFP output from Input A. Using untagged RFP gene in design III led to slight increase in overall RFP output but did not alleviate the signal balancing issue. The result implies that 5’ UTR structural effect is more dominant than RFP half-life in determining the success of layered XOR gate. In order to apply the XOR gate in the implementation of the half adder, design IV was characterised for its steady state profile by titrating with varying concentration of arabinose and rhamnose as shown in Fig. 5D. It is noteworthy that the XOR gate develop in this work possesses higher single cell computational capability as compared to that achieved by *Tamsir* and colleagues using a network of inter-communicating cells [38], hence circumventing problems associated with cell-cell communication.

### Design and Characterisation of Single Cell Half Adder and Half Subtractor

The half adder computes dual inputs with both AND and XOR logic operations to generate CARRY and SUM output, respectively. Building on bio-logical devices that were modularised and rigorously characterised earlier, we co-transformed constructs which produce GFP (CARRY) from HrpRS AND gate in low copy plasmid, RFP_asv_ (SUM) from hybrid promoters pBAD-Cl2B and pRHAB-Cl2B and λCl repressors from pHrpL promoter in high copy plasmid into *E. coli* (Fig. 6A). To study the digital performance of the single cell half adder, we characterised the system at both the population and single cell levels by microplate fluorescent assay (Fig. 6B) and flow cytometry (Fig. 6C, Supplementary Fig. 7) for four different logic conditions. The results show that the engineered cells exhibited robust and digital-like performance with minor expression leak (< 20%) in XOR output when both inputs were present. While previous characterisation with standalone XOR gates displayed near perfect XOR outputs, parallel implementation of both AND and XOR logic gates in half adder led to probable competition for HrpRS transcription activators by pHrpL promoters in both low and high copy plasmids - which is suggestive of expression shunting in competitive transcription dynamics [42]. In other words, the availability of HrpRS activators are divided between the pHrpL-GFP module in low copy plasmid and pHrpL-λCl module in high copy plasmid, thus causing both AND and XOR gates to perform below par compared to when they are operating individually. To affirm the hypothesis, we examined the AND output of standalone AND gate with the AND output of the half adder using microplate fluorescent assay. The results showed that the GFP output of isolated AND gate was approximately 7 times stronger than that of half adder’ s AND gate, thus confirming our hypothesis (Supplementary Fig. 8). It is noteworthy, that the reduced expression of GFP did not affect the overall performance of the half adder as effective half adder logic operations were still achieved. In the current single cell half adder, the engineered cells exhibited relatively healthy growth with the same order of viable cells (∼10^9^cfu/ml) in both induced and uninduced cell cultures (Supplementary Fig. 9). Nevertheless, as genetic complexity and heterologous expression increased, a concomitant increase in the metabolic burden in the *E. coli* cell was also observed.

**Fig. 7.**
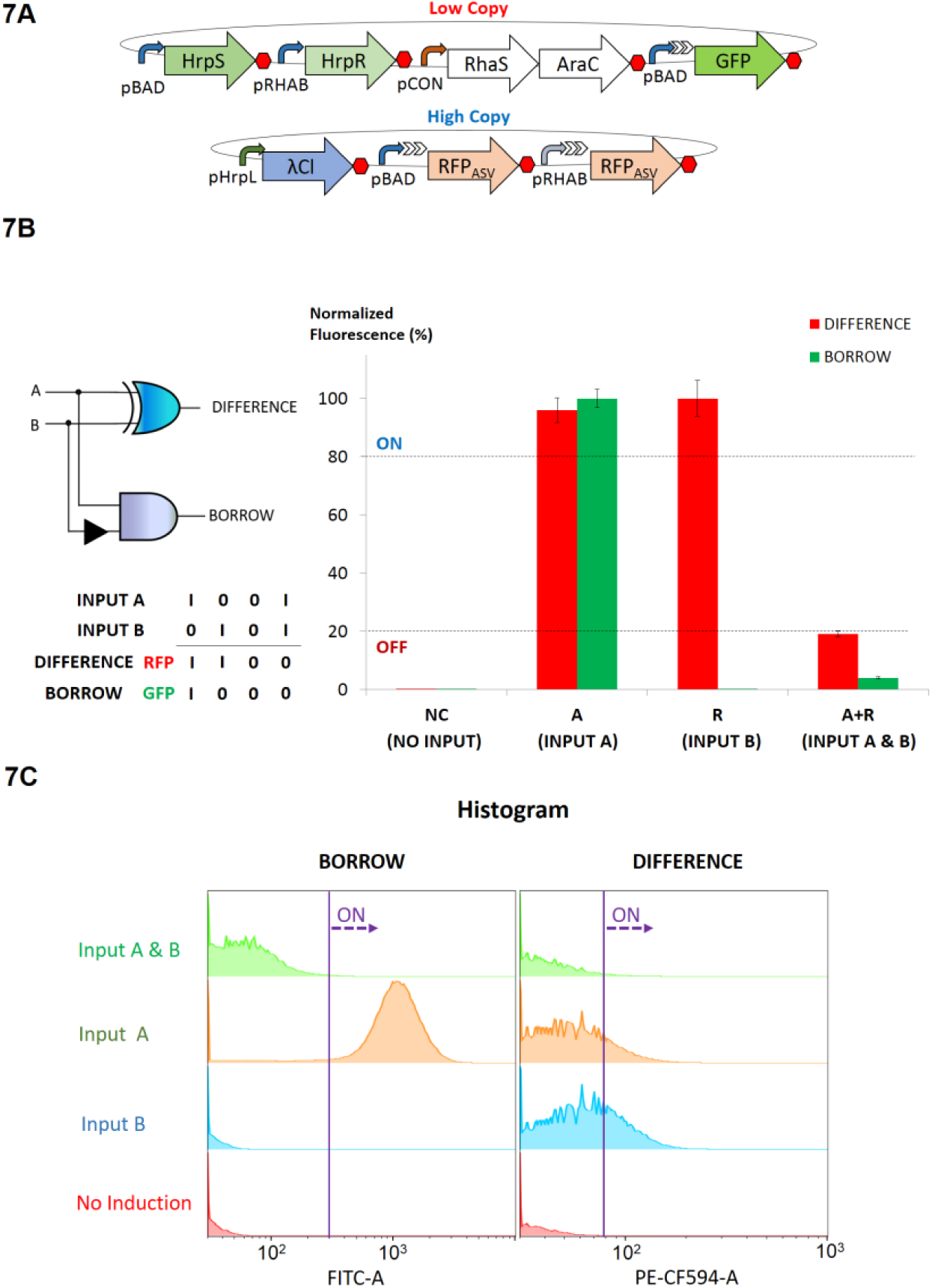
Design and characterization of the biological half subtractor.

A. Genetic blueprint of the half subtractor.
B. Digital performance of half subtractor at steady state.
C. Flow cytometry analysis of the half adder. Y axis coordinate represents population count, while FITC-A and PE-CF594-A represent channels that detects GFP and RFP fluorescence, respectively. Population shifts to the right represent ON behaviour. Error bars represent the standard deviation of four independent experiments.

To demonstrate the modularity of our approach, we also developed single cell half subtractor by performing slight modifications to the genetic circuits that formed the basis of the half adder. Specifically, GFP, which exemplifies BORROW output, was produced from the hybrid promoter pBAD-Cl2B in the low copy plasmid instead of the pHrpL promoter (Fig. 7A). As above, the construct which generated the BORROW output (GFP) and that which generated the DIFFERENCE output (RFP) were co-transformed into *E. coli* cells. Characterisation was undertaken at both the population and single cell levels by microplate fluorescent assay (Fig. 7B) and flow cytometry (Fig. 7C) under four different logic conditions. The results showed that the engineered cells functioned as effective biological half subtractors, producing GFP only in the presence of input A and RFP in the presence of input A or B, but not when both inputs were present.

## DISCUSSION

Logic gates are the basis of all electronic digital devices from mobile phones, to microprocessors, to computers. They are therefore the basis of the processing of information and control systems. Similarly, the development of biologically based logic gates and logical devices has major potential in terms of information processing and control. The design and testing of a half adder, which is the subject of this paper, is seen as a significant step in the development of biological logical devices, comprising multiple gates that work stably and in unison. Immediate areas of application are in advanced biosensors. In the longer term, there is the potential to development of biologically-based devices for information processing and control, for example in the application of human-imposed intracellular control. In the underlying strategy of the paper is one of applying systematic design through the application of engineering principles [43]. Using a forward engineering approach that is supported by modelling and rigorous characterisation, independent modules that enable programmable digital operations in prokaryotic cells, including simple genetic switches, AND, OR and NOT logic operations were systematically assembled and characterised. AND, OR and NOT logic gates were then layered in both parallel and serial arrangements to generate a repertoire of cellular Boolean operations that include NIMPLY, XOR, half adder and half subtractor logic operations. Using a bottom up approach for constructing biological systems of increasing complexity we assessed genetic architectures that led to genetic context dependent effects. On this basis, the significance of each design on the overall digital performance of programmable logic gates in engineered cells was studied, leading to the compilation of a comprehensive set of guidelines for troubleshooting synthetic genetic circuits (Table 1). This work together with recent studies conducted elsewhere, highlight the importance of modularity and characterisation during the systematic layering of multiple biological devices [10, 44, 45].

Overall, the presence of secondary structures in 5’-UTR of mRNA affects genetic expression most. We discovered that the presence of seven consecutive hairpins immediately downstream of promoter transcription start site would cause severe impediment of gene expression. Although OR gate design made up of tandem promoters can be subjected to the undesirable effects of 5’-UTR secondary structure, we showed that the effect is not pronounced in the digital performance of the OR logic when the promoters and DNA operator sites involved are of markedly different DNA sequences. The OR gate design that comprises a separate gene expression cassette also reliably demonstrates digital operation. However, the involvement of larger DNA modules and repetitive use of transcription terminators that are rich in secondary hairpin structures may impede system assembly in terms of construction efficiency and accuracy. Where identical DNA sequences are incorporated in a single mRNA transcript, as shown in a design II and III of XOR gate, the effect of 5’-UTR secondary structure preventing gene expression is significantly more pronounced. Thus, it is proposed that XOR gate logic in layered genetic circuits should be designed with two discrete expression cassettes instead of employing a tandem promoter circuit design. It would also be interesting to test if RNA processing tools can be employed in multiplex mode to insulate the myriad of biological devices from RNA genetic context dependent effects in layered genetic circuits concurrently.

Perhaps of particular interest, we discovered that σ54 promoters can exhibit genetic context dependent effects if two σ54 promoters are placed close to each other. Previously, σ54-dependent NtrC-binding promoters have been reported. These promoters permit transcription *in vitro* in the absence of enhancer-binding proteins and ATP under conditions that promote DNA melting. These include DNA supercoiling, temperature rise and lower ionic strength, or when characteristic point mutations are implemented on the σ54 protein [46, 47]. In this paper, we show that an upstream σ54 pHrpL promoter could also activate downstream pHrpL promoter *in vivo* if the two promoters are in close proximity - possibly as a result of plasmid DNA supercoiling. This undesired switched-on activity can be avoided by designing pHrpL expression modules in different plasmids, ie to use plasmid as genetic buffers to insulate such genetic context dependent effects.

While recombinases have been intelligently crafted into Boolean logic gates with DNA-encoded memory functions, it is important to note that biosensors connected in AND, OR and XOR operations with recombinase-based logic gates may not be able to distinguish inputs from different environments and provide the desired response. For example, a probiotic that is genetically programmed in AND logic to sense two inputs such as hypoxia and low pH may be activated for hypoxia and low pH signals in two different locations, as compared to sensing both signals *in situ*. The same may be applicable for other logic operations with recombinase-based logic gates. Thus, layered genetic circuits that are capable of sensing and providing location-sensitive Boolean logic operations are still useful in programming cellular behaviour. Of particular interest is a combination of layered genetic circuits, with the synthesis of recombinases as intermediary output, this may provide a novel and better platform for programmable cellular behaviour in terms of both accuracy and memory.

With the exceptions of a notable few [19, 48, 49], most studies of synthetic biological systems are centred on the development of rational engineering approaches, reporting successful and advantageous aspects of the engineered systems, with lesser focus on reporting failure modes and compiling the engineering solutions applied to troubleshoot system failures. As synthetic biology moves forward with greater focus on scaling the complexity of engineered genetic circuits, studies which thoroughly evaluate failure modes and engineering solutions will serve as important references for future design and development of synthetic biological systems.

## ACKNOWLEDGEMENT

We are grateful to P. Freemont, K. Jensen, C. Hirst and F. Jonas from Imperial College London for suggestions, M.W. Chang from the University of Singapore for experimental support and the use of flow cytometer for the work, and B.J. Wang for the kind donation of plasmids containing *hrpS*, *hrpR* and *pHrpL* promoter. We would also like to thank the anonymous reviewers for their comments on this paper.

